# Explainable HGT-based framework for predicting human dark kinase protein-pathway associations by leveraging BERT-based embeddings and WGAN-GP

**DOI:** 10.64898/2026.07.30.741784

**Authors:** Somarpita Dutta, Pralay Mitra

## Abstract

Discovery of pathway associations and druggability can leverage underutililized dark kinase genes for treating complex diseases (proven for cancer and neurodegeneration), boosted with computational methods. Herein, we employ BERT-based embeddings of proteins and pathways (refined via two-stage transformer and heterogeneous graph transformer) and protein-protein and protein-pathway associations-both positive (curated from databases) and negative (generated using Wasserstein Generative Adversarial Networks with gradient penalty) to train XGBoost and lightGBM classifiers for predicting pathways associated to human dark kinase proteins, with important features unveiled through SHAP analysis. All pathways are clustered and proteins related to same pathway clusters are grouped together (via predicted and positive protein-pathway associations). Selected PCOS-related human dark kinase proteins (with high predicted and existent associations to PCOS pathways) are docked with known PCOS drugs for druggability analysis. Our model attains accuracy, F1-score, specificity, MCC, AUROC and AUPRC of 0.9816, 0.9816, 0.9852, 0.9632, 0.9978 and 0.9982 respectively, supersedes existing work, correctly classifies 97.48% of test data, predicts 62225 pathway associations to above proteins, infers functional similarity of 96 such proteins to human protein(s) and traces nine important positive features. Our model can be used to determine varied functionalities and disease relevance of proteins via predicted pathway associations.

## 1 INTRODUCTION

Dark kinases, constituting of nearly one-third of the protein kinase gene family are largely understudied, poorly characterized and lack scientific annotations, but have recently shown promising results in treatment of cancer and neurodegeneration [1,2] and have been proved to be druggable [3]. Hence, harnessing capabilities of these dark kinases in treatment of other complex diseases, particularly with limited medications can boost their treatment.

Despite aforementioned efforts to highlight their potential, dearth of associations related to dark kinases is evident. They can be made utilisable in treatment of complex diseases with proper knowledge of pathway associations and drug interactions. RegPattern2Vec [4] aimed to predict pathway associations involving dark kinases. Biased random walks were performed to generate embeddings of dark kinases/pathways and logistic regression was considered to predict pathway associations of dark kinases. But, lack of associations pertaining to dark kinases built a poor network which also lessened the impact of network based methods. Reliance on structure, function and amino acid sequences of dark kinase proteins can ameliorate the situation. But they are poorly characterized, as mentioned earlier. Herein, language models have been found to be effective in capturing information on the above aspects based on protein sequences and provide fruitful predictions [5]. Since novel drugs require several years to be launched, repurposing existing drugs for treatment via dark kinase proteins as druggable targets can boost therapeutic strategies, particularly in complex diseases with multiple lines of treatment and those with limited medications like Polycystic Ovarian Syndrome (PCOS).

Here, we address the aforementioned gaps by predicting pathway associations of human dark kinase proteins (HDKPs) via XGBoost and LightGBM classifiers. Our training data comprises of modified BERT-based embeddings of proteins and pathways as well as protein-protein interactions (PPIs) and protein-pathway associations (PPAs). The BERT-based embeddings comprise of ESM2 embeddings [6](generated on protein sequences) for proteins and SBERT embeddings [7] (generated using pathway names) for pathways. Sets of ESM2 and SBERT embeddings, individually undergo the following transformations-reduction by PCA and Autoencoder (AE) [8], refinement through two-stage transformer-based encoder [9] and further reduction by PCA and AE. After undergoing these transformations, both embeddings sets are vertically stacked and further reduced by Heterogeneous graph transformer (HGT)[10]-based encoder. Positive (feasible) PPIs and PPAs consist of those procured from databases whereas WGAN-GP[11–13] is employed to generate negative (potentially infeasible) PPIs and PPAs. Both are included in training data. Our model attains accuracy, precision, recall, F1-score, specificity, MCC, AUROC and AUPRC (mean values) of 0.9816, 0.9852, 0.9780, 0.9816, 0.9852, 0.9632, 0.9978 and 0.9982 respectively, correctly classifies 97.48% of PPAs in test data and predicts 62225 PPAs. Also, it infers nine features which have pivotal and positive influence on predictions through SHAP analysis[14]. Combining predicted and positive PPAs (combined PPAs), we trace out number of associated PCOS pathways and percentage of total associated pathways related to PCOS of 57 PCOS-relevant HDKPs, sort them in descending order on these two aspects and analyse druggability of top ten (based on the sorting) by docking with nine well-known PCOS medicines. All ligand-protein combinations (best poses) are found to have binding energies in the range of -4.49 to -14.11 kcal/mol. Also, we group functionally similar proteins by conducting agglomerative hierarchical clustering on all pathways, mapping each pathway to its cluster label and each protein to a set of cluster labels corresponding to associated pathways (combined PPAs) and grouping proteins having same set of cluster labels. In this manner, we find out functional similarity of 96 HDKPs to one/more human proteins.

We contribute in the following ways-

- leverage BERT-based embeddings to predict pathway associations to HDKPs
- assess functional similarity of HDKPs to human proteins and thereby obtain a map of functionally similar proteins by dint of combined PPAs.
- analyse associations of HDKPs to PCOS by sorting them on the basis of number and percentage of PCOS pathways associated with respect to all associated pathways by utilizing combined PPAs.
- infer on druggability of top ten HDKPs based on sorting order through docking with well-known medications

Rest of the paper is structured as follows-we discuss the materials and methodology in section 2 followed by display and discussion of results in section 3, ending with conclusion in section 4.

## 2 MATERIALS AND METHODS

### 2.1 MATERIALS

Protein sequences of human proteins for generating ESM2 embeddings are curated from UniProt database [15]. Since, efficiency of ESM2 embeddings decreases beyond token size of 1024 [16], we restrict to 18054 human proteins. Pathway names of 2825 human pathways are curated from Reactome database [17] which will be used to generate SBERT embeddings. Data on parent-child relationship among above pathways is gathered from Reactome database.

PPAs, 103119 in number, are garnered from STRING [18] and CTD [19] databases. Out of 1477611 PPIs involving direct physical interactions between human proteins in STRING database, we consider 150931 PPIs with association scores equal to or higher than mean of the association scores of all PPIs. This is done to avoid overpopulating ID with PPIs. Proteins in STRING are denoted by ensembl [20] protein IDs which are mapped to UniProt IDs via UniProt mapping tool and proteins involved in one-to-one mapping of both IDs are considered above.

PCOS genes are accumulated from PCOSKB [21] and CTD databases. Genes are mapped to corresponding proteins (UniProt IDs) via UniProt mapping tool. Gene-protein pairs having one-to-one mapping are retained and proteins in these pairs constitute of PCOS proteins in our study. Pathways related to PCOS are gathered from CTD. Files denoting two-dimensional structure (.sdf) of nine PCOS medicines are procured from DrugBank database [22]. Altogether, 16599 proteins and 738 pathways related to PCOS are curated.

Dark kinases in Dark Kinase Knowledgebase [23] are mapped to human proteins via UniProt mapping tool. Adhering to one-to-one mapping, 129 HDKPs are found in human proteins in our study and considered here.

### 2.2 METHODS

#### 2.2.1 PROCEDURAL STEPS

Refined Feature Vector (FV) set generation: Embeddings of size 1280 for human proteins are generated by ESM2 considering protein sequences as input. SBERT is used to generate 384-dimensional embeddings for pathways based on pathway names. ESM2 embeddings are reduced by PCA then AE, refined through two-stage transformer encoder stack and again reduced by PCA and AE to generate reduced protein FV set. Similar transformations are applied on SBERT embeddings to generate reduced pathway FV set. Each FV in reduced protein and reduced pathway FV set is of size 32. FVs of these two FV sets are vertically stacked to generate latent FV set having 20879 (18054 proteins+2825 pathways) FVs (embeddings), each of size 32. The latent FV set is finally refined by HGT-based encoder (which considers latent FV set and interaction data as input) to produce the refined FV set.

Interaction Data (ID) and test data generation: ID consists of PPIs and PPAs. Positive ID, accounting for feasible interactions/associations consist of whole of curated PPIs and around 80% of curated. PPAs (20% reserved for test data). Potentially infeasible PPIs and PPAs constitute of negative ID. These are generated via WGAN-GP which considers the positive ID and latent FV set as input. ID consists of positive ID and negative ID. Positive ID has 150931 PPIs and 82495 PPAs and negative ID possesses 170224 PPIs and 55826 PPAs. Altogether, ID has 464107 interactions/associations in its repertoire. Test data has 20599 feasible (positive) PPAs after removing those present in ID.

Blind data generation: Based on pathway tree constructed from parent-child relationship (curated from Reactome database), 663 pathways are inferred as leaf nodes. An exhaustive combination of human dark kinase proteins and above pathways, after removing PPAs in both ID and test data, deduces to 85392 links in blind data.

Training and prediction using lightGBM and XGBoost classifiers: Proteins/proteins and pathways serve to be interacting/associated entities in PPIs/PPAs respectively. For each interaction/association in ID, FVs of interacting entities from refined FV set are concatenated and fed to both lightGBM (implemented using lightgbm [24]) and XGBoost (implemented via XGBoost [25]) classifiers. Both classifiers predict probabilities of feasibility for blind and test data associations. Considering 0.7 as threshold for predicting feasible associations, both classifiers together classify 97.48% of test data correctly and predict 62225 PPAs in blind data. Generation of refined FV set, ID, test data and blind data is shown in figure 1.

**Figure 1.**
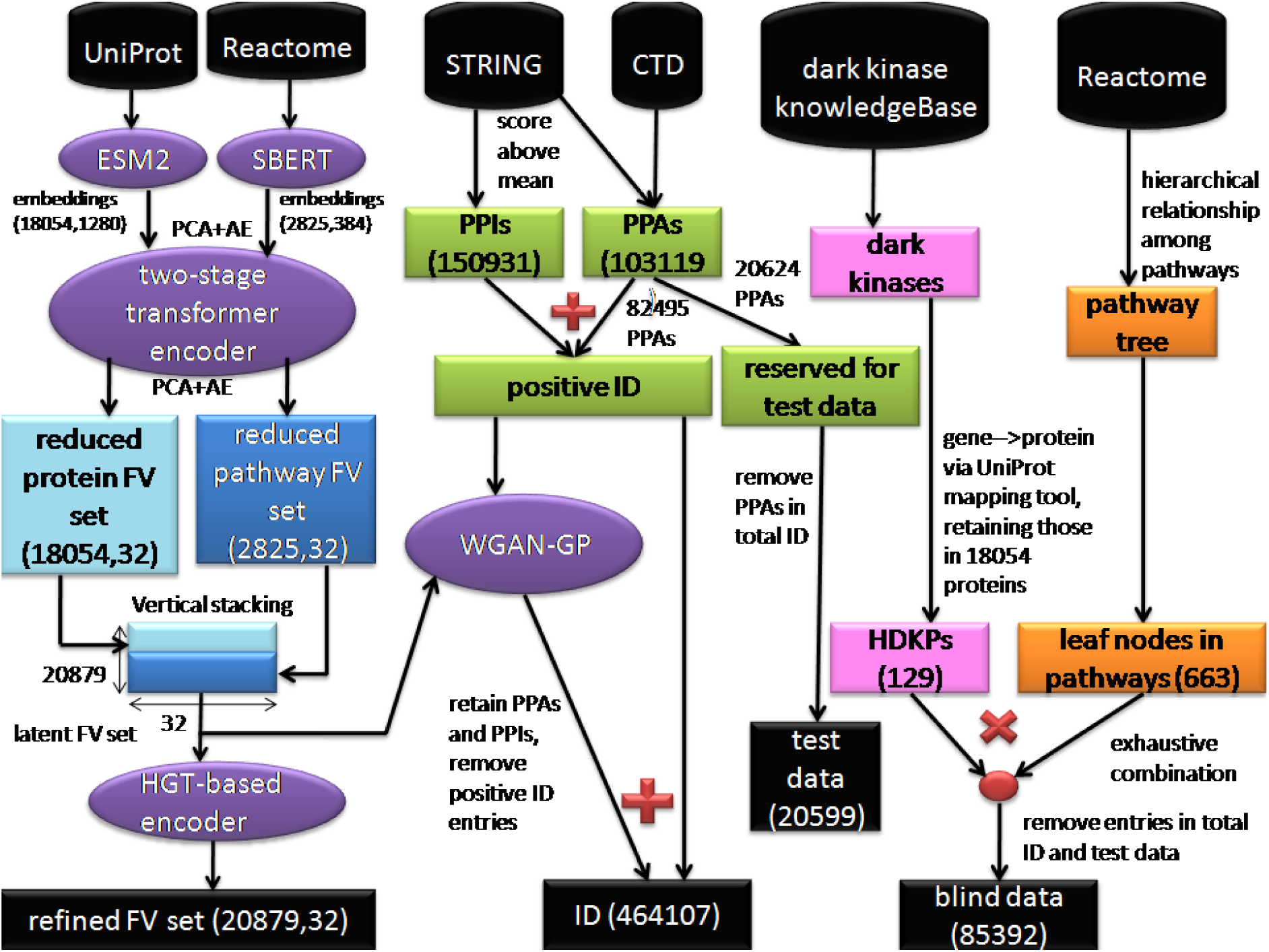
construction of refined FV set, ID, test data and blind data. ESM2 embeddings of size 1280 (on 18054 human proteins) and SBERT embeddings of 384 dimensions (on 2825 pathways) are independently reduced by PCA then AE, refined through two-stage transformer encoder and further reduced via PCA and AE to generate reduced protein FV set and reduced pathway FV set respectively. Both these FV sets are vertically stacked to construct latent FV set, which is passed through HGT-based encoder to produce refined FV set having 20879 FVs (18054 human proteins and 2825 pathways), each of size 32. The latent FV set is inputted to WGAN-GP along with positive ID (made up of 82495 PPAs curated STRING and CTD and 150931 PPIs curated from STRING) to generate negative ID. Both positive and negative ID combine to produce ID. Out of 103119 PPAs curated in total, 20624 are reserved for test data, and after removing entries in ID, test data consists of 20599 PPAs. Dark kinases are curated from dark kinase knowledgebase, which are converted to HDKPs via UniProt mapping tool (through one-to-one mapping between genes and proteins). A pathway tree is built out of hierarchical relationship between human pathways in Reactome database, and the leaf human pathways are pointed out. An exhaustive combination of HDKPs and the above pathways, after removing ID and test data entries, contained 85392 entries.

SHAP analysis: Features of proteins and pathways which play pivotal roles in model (XGBoost and LightGBM) predictions are inferred through SHAP (SHapley Additive exPlanations) analysis on test data (separately for both classifiers). For each feature, we express its importance across the test data via normalized SHAP weights (values) (dividing the mean absolute SHAP value of that feature by sum of the mean absolute SHAP values of all other features) and percentage-wise normalized SHAP weights (scaling the normalized SHAP values to 100%). If increasing/decreasing the value of a feature improves predictions of our model, that feature is assumed to exhibit positive/negative influence on the model performance. This is revealed through beeswarm plot for top 20 important features (based on decreasing values of percentage-wise SHAP weights). In all, ten and nine protein and pathway features are considered the most important, out of which six and three are found to exhibit positive influence on model performance.

PCOS relevance and druggability: A total of 57 HDKPs are found to be included in PCOS proteins (curation discussed under ‘Materials’ in section 2.1), hence inferred as PCOS relevant. These HDKPs are further sorted based on association with number of PCOS pathways and their ratio with respect to total pathways associated, expressed as percentage. It is noteworthy that both predicted PPAs and PPAs in positive ID are considered here. This is portrayed in figure 2a. The ones in top ten are considered for further analysis of druggability by docking with nine PCOS drugs. Detailed discussion is provided in in section 3.6 under ‘PCOS association and druggability analysis’.

**Figure 2.**
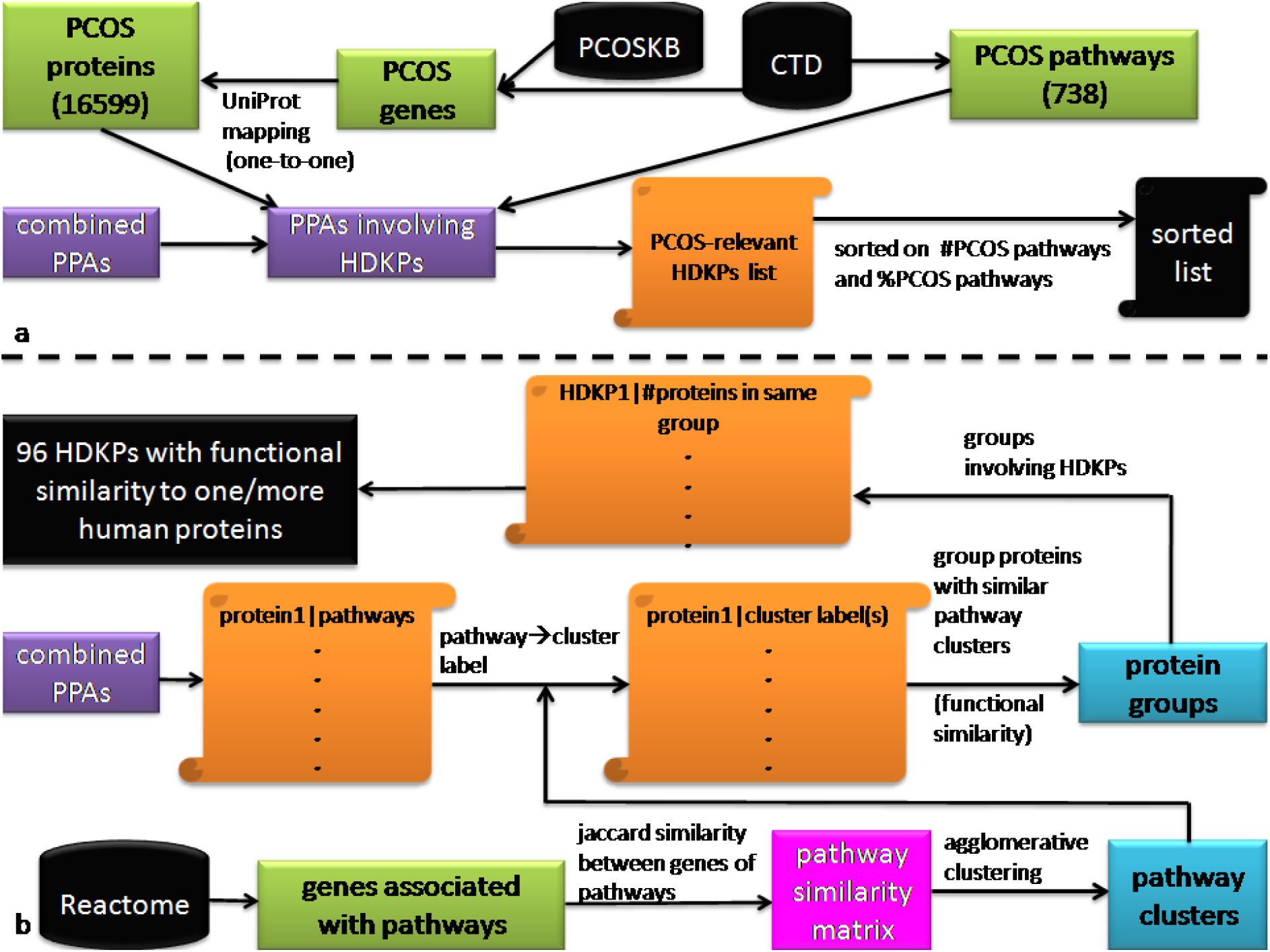
a Construction of PCOS-associated pathways of PCOS-relevant HDKPs. PPAs involving PCOS-relevant HDKPs (based on intersection of HDKPs and PCOS proteins mapped via UniProt mapping tool, from PCOS genes, curated from PCOSKB and STRING database) are transformed into a list containing number and percentage of PCOS-associated pathways for each PCOS-relevant HDKP, sorted further on these two parameters. Figure 2b Functional similarity of HDKPs to one/more human proteins. Similarity between pathways is estimated via Jaccard similarity between genes associated with pathways, generating pathway similarity matrix. This is inputted to agglomerative clustering to generate pathway clusters. Each protein-pathway association is transformed to protein-cluster(s) (cluster(s) in which the pathway is included) association. All cluster labels corresponding to each protein are considered separately and proteins associated with same cluster labels are grouped together, resulting in 96 HDKPs as functionally similar to one/more human proteins.

Clustering analysis: All pathways in training data are assessed for mutual similarity by dint of Jaccard similarity between genes associated with corresponding pathways. A pathway similarity matrix is built out of these similarity values which is used to cluster pathways through hierarchical agglomerative clustering. Fifty uniformly-spaced values in the range of 0.1 to 0.5 are considered as distance threshold in clustering for deciding on optimal clusters and optimal distance threshold, deduced using Within Cluster Sum-of-Squares (WCSS) strategy combined with elbow method (implemented using kneed library[26]). PPAs (predicted and positive) are used to map all proteins in training data to pathway clusters representing pathways associated with respective proteins. Proteins having similar pathway clusters are grouped together. In this way, all proteins are grouped based on clustering of associated pathways. Proteins in same group are considered functionally similar. We consider groups containing HDKPs and find 96 HDKPs similar to one/more proteins and 33 with none. This is pictorially shown in figure 2b. Data on functional similarity of HDKPs to human proteins is available in supplementary table 1. The whole process of implementation is shown in figure 3.

**Figure 3.**
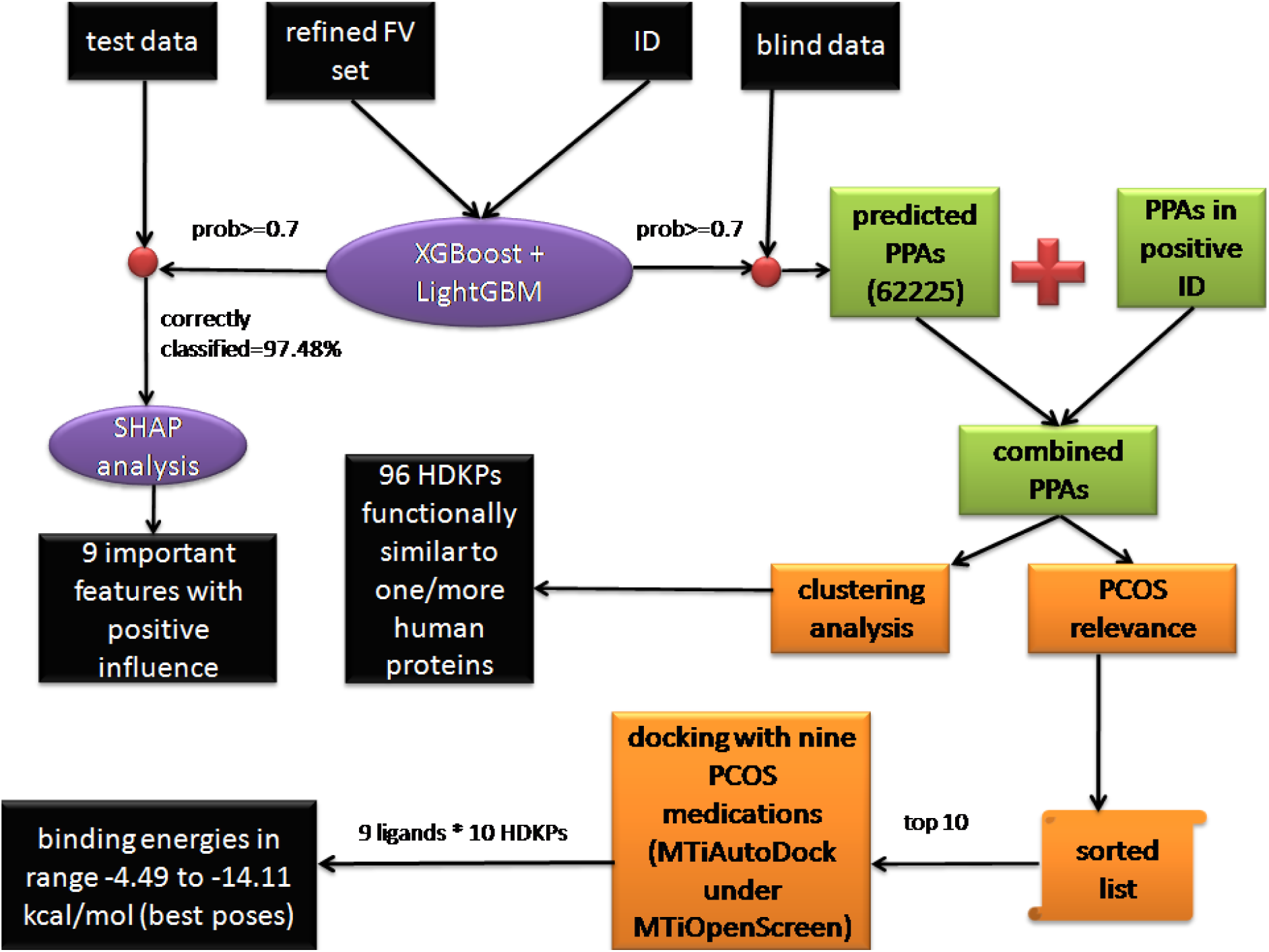
Overall flow of implementation. The refined FV set and ID are used to train XGBoost and LightGBM classifiers. All predictions with a probability of 0.7 or more are considered feasible. 9 important features having positive influence on predictions are traced out through SHAP analysis on test data. Predicted PPAs are combined with PPAs in positive ID to generate combined PPAs. Using these combined PPAs, on one hand, we conduct clustering analysis (shown in figure 2b), outputting 96 HDKPs which are functionally similar to one/more human proteins. On the other hand, we assess PCOS relevance of HDKPs (shown in figure 2a), which results in a list of PCOS-relevant HDKPs, sorted as per their number and percentage of associations. Considering top ten among those in sorted list, we perform docking with nine PCOS medications via MTiAutoDock service of MTiOpenScreen web server. Considering best poses, binding energies of all ligand-protein combinations are in the range of -4.49 to -14.11 kcal/mol.

#### 2.2.2 ELEMENTS OF FRAMEWORK

Here, we will discuss four important components of our framework, namely vanilla autoencoder, two-stage transformer-based encoder, WGAN-GP and HGT-based encoder. For each of these components, we tune the hyperparameters through 50 trials of Optuna-based hyperparameter tuning [27] strategy. This helps us in getting optimal hyperparameters which ensure optimal performance of the framework components. Also, in this section, for convenience in discussion, we denote proteins/pathways as nodes and interactions/associations as edges.

Vanilla Autoencoder: It has three main components-encoder, bottleneck (latent space) and decoder. The encoder consists of two stages-input projection and feature refinement. The input projection stage consists of linear projection layer to project data into lower-dimensional space followed by ReLU layer. The output of this stage is fed to feature refinement stage which reduces the dimension further. The output of this stage is passed through the bottleneck (latent space) which finally reduces the input to desired reduced dimension. The decoder consists of two stages for reconstruction of original input, each consisting of a linear layer followed by ReLU activation layer. This is followed by the final output stage consisting of a linear layer followed by sigmoid activation layer. The hyperparameters include final reduced dimension of embeddings, intermediate reduced dimension of embeddings, learning rate, batch size and number of epochs.

Two-stage transformer-based encoder: It majorly consists of two transformer blocks. Each block consists of the following stages-first layer normalization, multi-head self-attention (MHSA), first dropout, first residual connection, second layer normalization, feedforward neural network (FFN), second dropout and second residual connection. Our data is inputted to the first transformer block. The output of the first block serves as input to the second block. The output of the second block denotes refined context-aware embeddings. Layer normalization is performed to prevent exploding gradients. MHSA stage performs three internal projections for every embedding-query (Q), key (K) and value (V). Each embedding is split into multiple heads-each focussing on a separate relationship. FFN, consisting of two layers and GELU activation function extracts deeper patterns within embeddings by expanding them four times and shrinking back. Residual connections enable preservation of original data along with learning the refinements. Addition of dropout layers prevent overfitting of data. The hyperparameters considered here include number of heads for MHSA, dropout and learning rate. The architecture is inspired from Vaswani et al[9].

WGAN-GP: It consists of a generator and a discriminator. The generator aims to create fake edges (interactions/associations) and discriminator aims to recognise them. Repeating this process multiple times ensures generation of edges which are hard to discriminate from positive ID. The generator consists of a linear layer for projecting noise into higher dimensional space, a ReLU activation layer and another linear layer for mapping features to a pair of values. These pair of values are passed through sigmoid activation layer, multiplied by total nodes (proteins/pathways)-1, rounded and clamped to generate node indices as potential hard negative samples. The discriminator, similarly consists of a linear layer to project the batch of edges to higher dimensional space and reduce them to hidden dimension, followed by ReLU activation layer and another linear layer for projecting the reduced embeddings to a critic score for each edge in the batch. Training process involves training generator 100 times and each time the generator is trained, the discriminator is trained five times prior to generator. A sequence of events take place during each training of discriminator. Firstly, the discriminator (critic) considers a batch of edges from positive ID, concatenates embeddings of nodes representing these edges and scores these edges. Secondly, the generator suggests a set of node pairs based on random noise and concatenates embeddings of representative nodes like discriminator. Thirdly, interpolated edges are created by rounding edges created by both generator and discriminator and the gradient of the discriminator output is calculated with respect to these interpolated edges. After the training of discriminator, the generator creates fake node pairs, inputs them to discriminator and updates itself based on discriminator score. Based on repeated training and updation, the discriminator score improves and the negative of this score (considered as generator loss) is used for concluding on optimal hyperparameters. WGAN-GP is trained once more on the optimal hyperparameters concluded to produce the final trained generator. This generator goes in evaluation mode to generate hard negative samples equal in number to the positive edges. The hyperparameters include learning rate of generator, learning rate of discriminator, batch size, hidden dimension of generator and hidden dimension of discriminator. The architecture of WGAN-GP is based on the work of Argovsky et al.[13]

HGT-based encoder: It mainly consists of a linear projection layer followed by HGTConv layers (implemented using Pytorch Geometric [28]) along with normalization and ReLU activation layers and a final output linear layer. Alongside, there is link prediction function which calculates dot product of node embeddings involved in binary interactions and outputs score as well as training and validation loss during each trial of optuna-based hyperparameter tuning. The latent FV set and ID are inputted to the linear projection layer. This layer projects the embeddings onto a shared space (specified by dimension of refined embeddings) and ensures uniform length of embeddings for all nodes. The output of linear projection layer is passed through HGTConv layers which employ multi-head attention to focus on related interactions where attention weights are specific to relationship types. Each HGTConv layer is followed by normalization and ReLU activation layers to generate scaled and non-linear features. The number of HGTConv layers are inferred through optuna-based hyperparameter tuning. The learned features from the final HGTConv block are passed through a linear layer to generate refined embeddings. The hyperparameters considered include number of attention heads, number of HGTConv layers and learning rate. We have set the dimension of refined embeddings to size of embeddings in latent FV set. This is because we want to refine, and not reduce these embeddings further. The architecture is developed closely in lines of official implementation of heterogeneous graph transformer [10] and Pytorch Geometric documentation.

XGBoost and LightGBM classifiers: For both of these classifiers, tuning of hyperparameters is performed first. Based on the ideal hyperparameters inferred, the classifier is trained once again followed by classification of test data and prediction on blind data. We consider hyperparameters like learning_rate, maximum number of levels of growth of each decision tree (DT) (max_depth), number of DTs (n_estimators), fraction of training data for training each DT (subsample), fraction of features (columns) for constructing each DT (colsample_bytree), regularisation parameters to prevent overfitting (gamma, reg_alpha, reg_lambda) and minimum sum of instance weights for creating a new node (min_child_weight) for training XGBoost. Similarly, for lightGBM, we consider hyperparameters like n_estimators, learning_rate, maximum possible number of leaves in each DT (no_of_leaves), max_depth, minimum data samples to create new leaf node (minimum_child_samples), subsample, colsample_bytree, reg_alpha and reg_lambda. Optimized values of hyperparameters for all components of the framework are available in supplementary table 2.

## 3 RESULTS AND DISCUSSION

### 3.1 PERFORMANCE OF OUR MODEL

Here, we assess the performance of our model in 10 iterations of stratified 10-fold cross validation using mean values of eight performance metrics namely accuracy, precision, recall, F1-score, specificity, MCC, AUROC and AUPRC across all folds and iterations. Although, accuracy focuses on number of correct predictions, AUROC and AUPRC provide better estimates for skewed datasets. High values of precision and recall point at reduced false positives and false negatives. F1-score, a balance of both, keeps both false positives and false negatives at bay. Specificity determines identification of actual negatives. Last, but not the least, MCC determines overall status of model performance and is effective for imbalanced data.

Here, we evaluate model performance in three different setups. In the first case, the refined FV set and ID are inputted to XGBoost and LightGBM classifiers. If we consider a network consisting of 18054 proteins and 2825 pathways as nodes and entries in ID as edges between nodes, we find out 3229 isolated nodes and the largest connected component having 17625 nodes. Accordingly, in the second case, we remove isolated proteins and pathways from refined FV set and retrain both classifiers and evaluate performance. In the third case, we retain only proteins and pathways in the largest connected component in the refined FV set. ID is also modified to contain interactions/associations between the aforementioned nodes. Both classifiers are trained on this combination and the performance is assessed.

The performance in all three cases is enumerated in table 1. In all three cases, the performance of both classifiers is denoted by the mean values with standard deviation in brackets for all of the performance metrics. The performance of our model is denoted as the mean of mean values reported for both classifiers for all performance metrics with standard deviation in brackets. The mean values of performance metrics remain almost similar in all three cases with difference in standard deviation. The difference in performance metrics between lightGBM and XGBoost classifiers gradually lessens as we move from cases first to third, leading to decrease in standard deviation for all performance metrics of our model. Since 18 HDKPs exist as isolated nodes, we continue with refined FV set and ID as training data for further stages of assessment.

**Table 1.** Table showing performance of our model on whole training data, after removing isolated nodes and considering the strongest connected component.

| performance metric | LightGBM<br>(±standard deviation) | XGBoost<br>(±standard deviation) | Our model<br>(±standard deviation) | LightGBM<br>(±standard deviation) | XGBoost<br>(±standard deviation) | Our model<br>(±standard deviation) | LightGBM<br>(±standard deviation) | XGBoost<br>(±standard deviation) | Our model<br>(±standard deviation) |
| --- | --- | --- | --- | --- | --- | --- | --- | --- | --- |
|  | whole training data |  |  | removing isolated nodes |  |  | considering strongest connected component |  |  |
| accuracy | 0.9819(± 0.0005) | 0.9812(± 0.0006) | 0.9816(± 0.0004) | 0.9819(± 0.0005) | 0.9817(± 0.0005) | 0.9818(± 0.0001) | 0.9819(± 0.0006) | 0.9818(± 0.0006) | 0.9818(± 0.0) |

| AUPRC | AUROC | MCC | specificity | f1_score | recall | precision |
| --- | --- | --- | --- | --- | --- | --- |
| 0.9982(±<br>0.0001) | 0.9979(±<br>0.0001) | 0.9639(±<br>0.0011) | 0.9859(±0<br>.0009) | 0.982(±0<br>.0005) | 0.9781(±<br>0.001) | 0.9859(±<br>0.0009) |
| 0.9981(±<br>0.0001) | 0.9978(±<br>0.0001) | 0.9625(±<br>0.0011) | 0.9845(±0<br>.0008) | 0.9813(±<br>0.0006) | 0.978(±0<br>.001) | 0.9846(±<br>0.0008) |
| 0.9982(±<br>0.0) | 0.9978(±<br>0.0) | 0.9632(±<br>0.0007) | 0.9852(±0<br>.0007) | 0.9816(±<br>0.0004) | 0.978(±0<br>.0) | 0.9852(±<br>0.0006) |
| 0.9981(±<br>0.0001) | 0.9979(±<br>0.0001) | 0.9638(±<br>0.0011) | 0.9857(±0<br>.0008) | 0.9819(±<br>0.0005) | 0.9782(±<br>0.0009) | 0.9857(±<br>0.0008) |
| 0.9981(±<br>0.0001) | 0.9979(±<br>0.0001) | 0.9634(±<br>0.0011) | 0.9854(±0<br>.0009) | 0.9817(±<br>0.0005) | 0.978(±0<br>.0009) | 0.9855(±<br>0.0008) |
| 0.9981(±<br>0.0) | 0.9979(±<br>0.0) | 0.9636(±<br>0.0002) | 0.9856(±0<br>.0001) | 0.9818(±<br>0.0001) | 0.9781(±<br>0.0001) | 0.9856(±<br>0.0001) |
| 0.9981(±<br>0.0001) | 0.9979(±<br>0.0001) | 0.9638(±<br>0.0013) | 0.9856(±0<br>.0007) | 0.9819(±<br>0.0006) | 0.9782(±<br>0.0009) | 0.9856(±<br>0.0007) |
| 0.9981(±<br>0.0001) | 0.9979(±<br>0.0001) | 0.9636(±<br>0.0012) | 0.9853(±0<br>.0007) | 0.9818(±<br>0.0006) | 0.9783(±<br>0.0009) | 0.9854(±<br>0.0007) |
| 0.9981(±<br>0.0) | 0.9979(±<br>0.0) | 0.9637(±<br>0.0001) | 0.9854(±0<br>.0002) | 0.9818(±<br>0.0) | 0.9782(±<br>0.0) | 0.9855(±<br>0.0001) |

### 3.2 COMPARISON WITH EXISTING WORK

Here, we compare our study with an existing work, RegPattern2Vec [4]. It uses biased random walks to generate embeddings and uses logistic regression as classifier for predicting pathway associations of dark kinases. Since, RegPattern2Vec uses F1-score and AUROC to benchmark performance, we adhere to these two metrics here and the performance of both is shown in figure 4. The performance of both is noted on logistic regression classifier. We supersede RegPattern2Vec in both metrics which speaks of the superiority of our approach. Additional information regarding this comparison are available in supplementary table 5.

**Figure 4.**
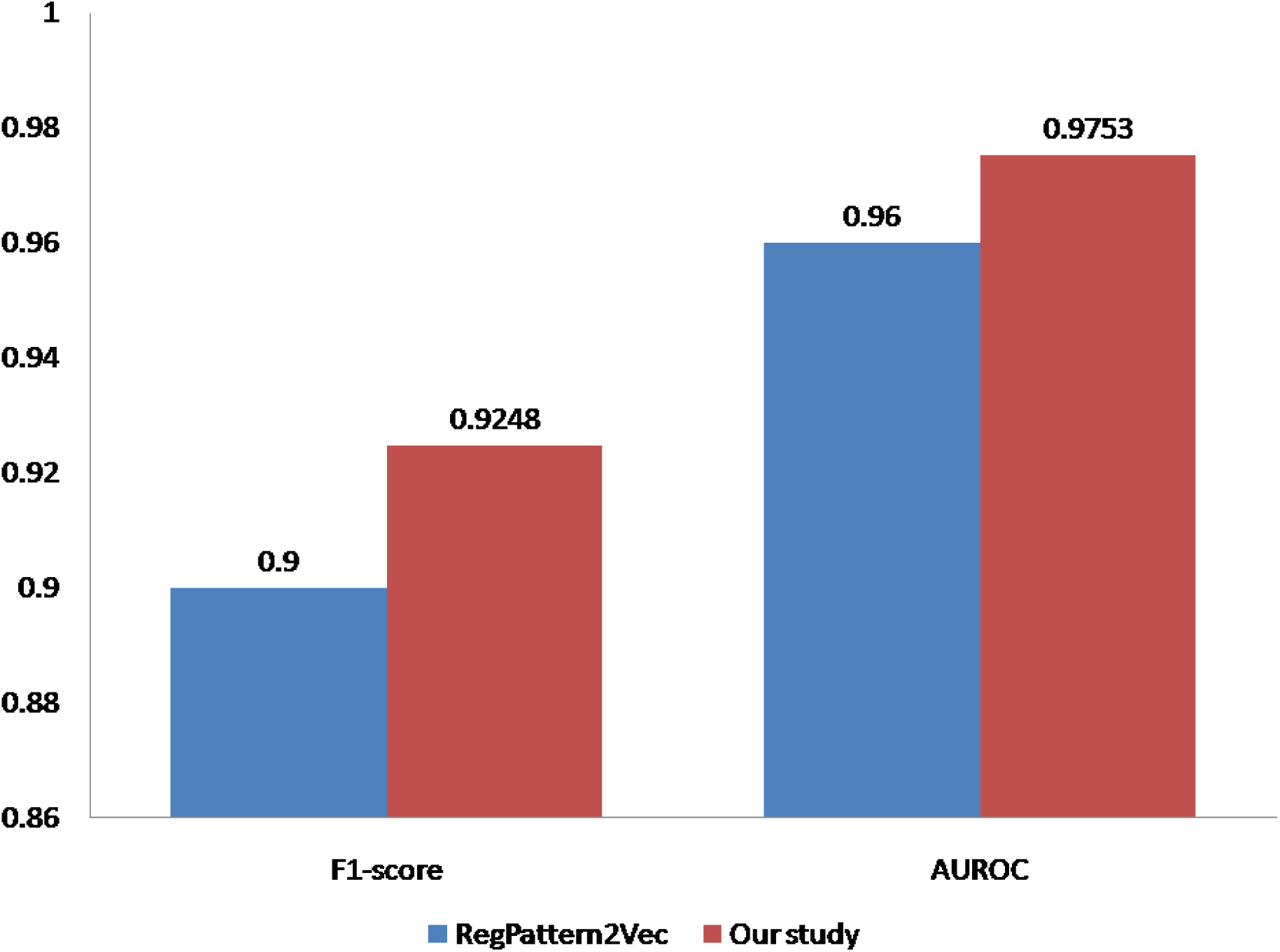
Comparison of performance of our model with RegPattern2Vec, both using logistic regression as classifier.

RegPattern2Vec serves to be a pioneering work in predicting associations between pathways and dark kinases. We move a step ahead in analysing the utility of HDKPs in PCOS associations by focusing on associations with PCOS pathways and druggability through docking with popular drugs for PCOS treatment (anovulation, insulin resistance and hyperandrogenism). Also, we attempt to find functional similarity of HDKPs to other human proteins through clustering of associated pathways.

### 3.3 PREDICTION ON TEST DATA AND BLIND DATA

As shown in figure 5, XGBoost has a better performance in test data validation with 20165 PPAs correctly classified in comparison to 20161 PPAs in lightGBM. Also, XGBoost predicts 64239 PPAs whereas lightGBM predicts 64354 PPAs.

**Figure 5.**
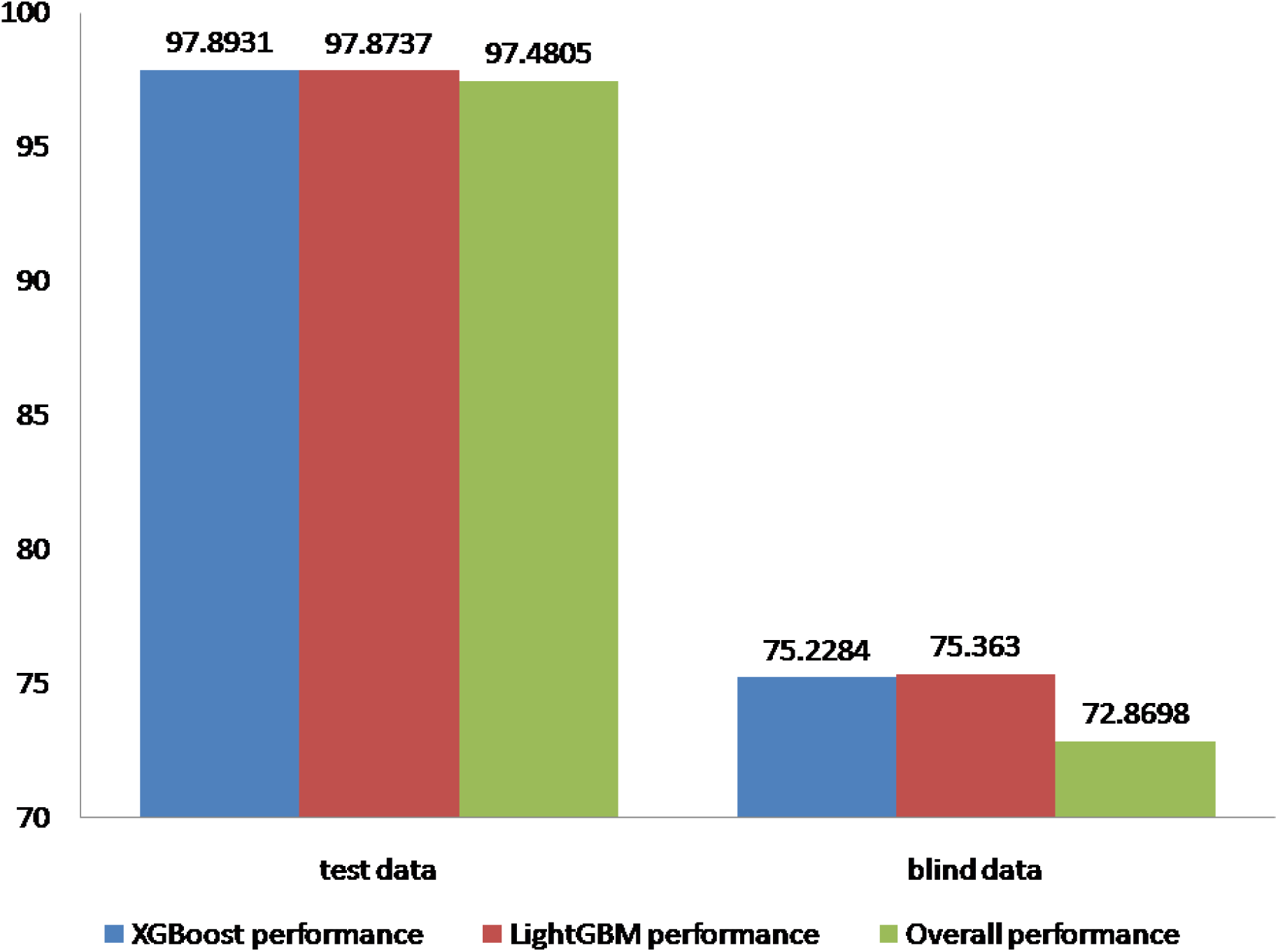
Comparison of performance of XGBoost and lightGBM classifiers on test data and blind data. The combined prediction of both classifiers is also expressed alongwith.

### 3.4 ANALYSIS ON FEATURE IMPORTANCE: SHAP ANALYSIS

We conduct SHAP analysis on test data for both XGBoost and LightGBM classifiers features of FV which are paramount to model performance. Features of proteins are identified as 1 to 32 and pathways as 33 to 64. There are two points to consider while assessing influence of features on model performance – how much the impact is pronounced and whether the influence is positive or negative. Relative importance of features, expressed as SHAP weights and percentage-wise SHAP weights is enumerated in table 2. Nature of influence is observed through beeswarm plot shown in figure 6 for XGBoost. The same is observed in figure 7 for lightGBM. In XGBoost, 10 proteins and 10 pathway features fall in top 20 most influential features. Out of these, six and three are found to exhibit positive influence. Similarly, in lightGBM, the most significant features include 11 from proteins and nine from pathways, with six protein-based and three pathway-based features having positive influence on model performance. SHAP weights and percentage-wise SHAP weights for all features are available in supplementary tables 3 (XGBoost) and 4 (lightGBM).

**Figure 6.**
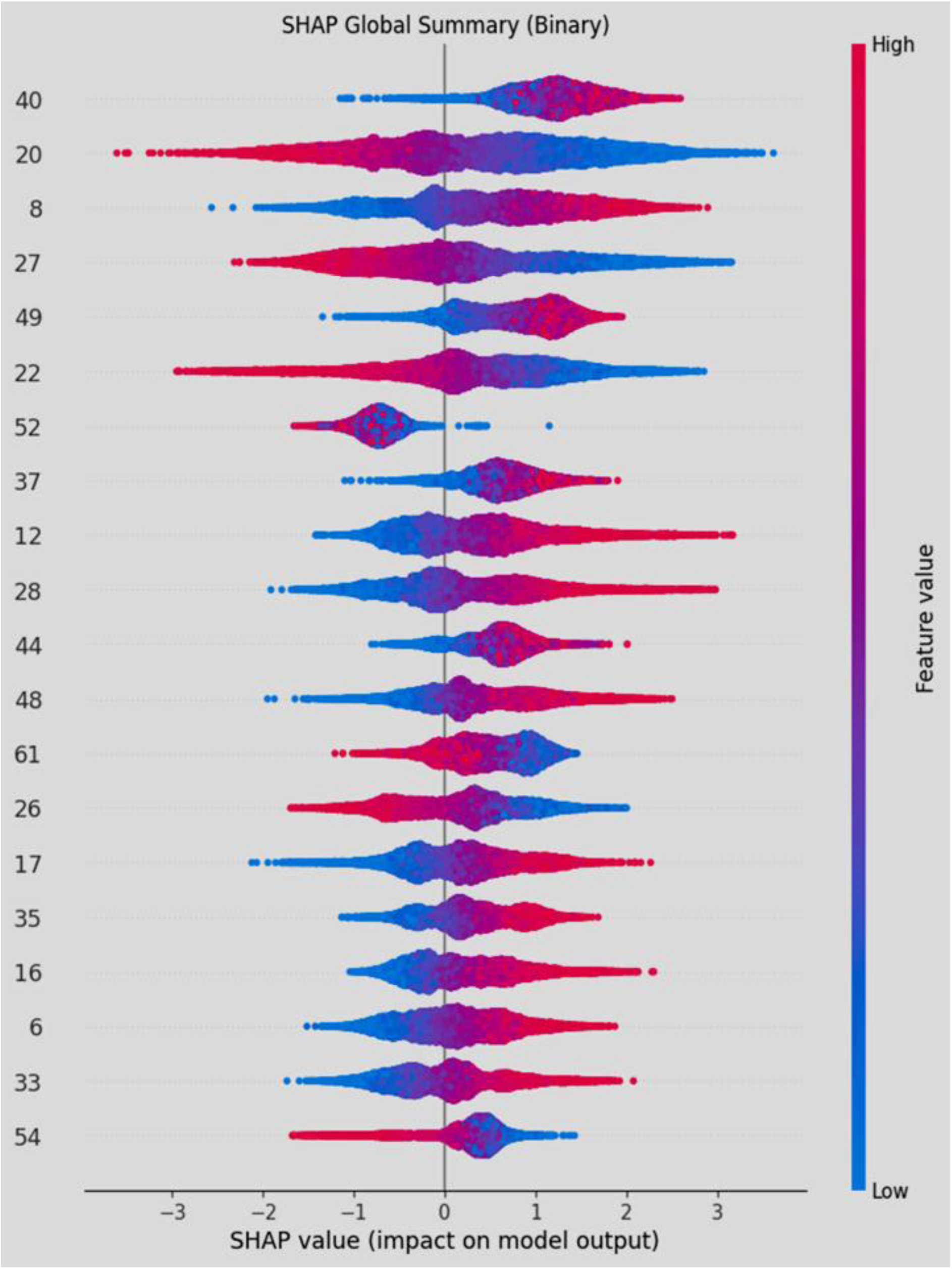
SHAP analysis on XGBoost classifier.

**Figure 7.**
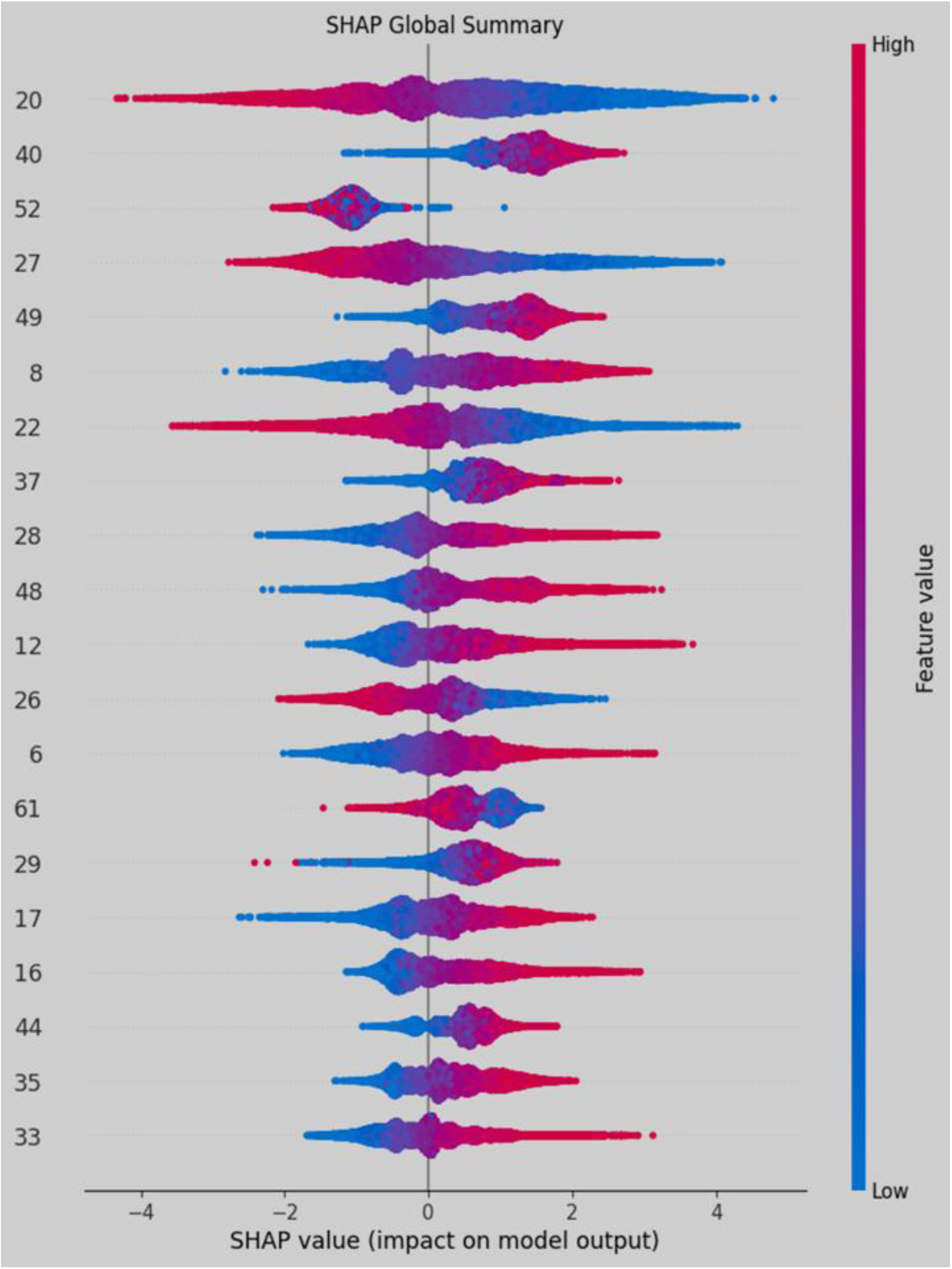
SHAP analysis on LightGBM classifier.

**Table 2.** Table showing top 20 features as per SHAP analysis on XGBoost and LightGBM classifiers.

| XGBoost classifier SHAP analysis | LightGBM classifier SHAP analysis |
| --- | --- |

| *feature* | *Protein*<br>*/pathway* | *SHAP*<br>*weights* | *Percentage*<br>*-wise*<br>*SHAP*<br>*weights* | *feature* | *protein/pat*<br>*hway* | *SHAP*<br>*weights* | *Percentage*<br>*-wise*<br>*SHAP*<br>*weights* |
| --- | --- | --- | --- | --- | --- | --- | --- |
| feature:40 | pathway | 1.193298 | 5.851591 | feature:20 | protein | 1.326707 | 5.652111 |
| feature:20 | protein | 1.005577 | 4.931058 | feature:40 | pathway | 1.280533 | 5.455398 |
| feature:8 | protein | 0.826984 | 4.055291 | feature:52 | pathway | 1.113551 | 4.744013 |
| feature:27 | protein | 0.797848 | 3.912416 | feature:27 | protein | 0.966967 | 4.119527 |
| feature:49 | pathway | 0.77147 | 3.783064 | feature:49 | pathway | 0.949602 | 4.04555 |
| feature:22 | protein | 0.76028 | 3.728196 | feature:8 | protein | 0.896918 | 3.821101 |
| feature:52 | pathway | 0.73441 | 3.601336 | feature:22 | protein | 0.891685 | 3.798809 |
| feature:37 | pathway | 0.630069 | 3.089679 | feature:37 | pathway | 0.725049 | 3.088894 |
| feature:12 | protein | 0.611639 | 2.9993 | feature:28 | protein | 0.692992 | 2.952323 |
| feature:28 | protein | 0.600424 | 2.944306 | feature:48 | pathway | 0.687569 | 2.929219 |
| feature:44 | pathway | 0.582893 | 2.858341 | feature:12 | protein | 0.619899 | 2.64093 |
| feature:48 | pathway | 0.572985 | 2.809754 | feature:26 | protein | 0.612735 | 2.610409 |
| feature:61 | pathway | 0.544915 | 2.672108 | feature:6 | protein | 0.600846 | 2.559757 |
| feature:26 | protein | 0.541711 | 2.656395 | feature:61 | pathway | 0.585319 | 2.493608 |
| feature:17 | protein | 0.486247 | 2.384415 | feature:29 | protein | 0.584374 | 2.489582 |
| feature:35 | pathway | 0.460145 | 2.256419 | feature:17 | protein | 0.577987 | 2.462372 |
| feature:16 | protein | 0.456338 | 2.23775 | feature:16 | protein | 0.561964 | 2.394111 |
| feature:6 | protein | 0.455914 | 2.235671 | feature:44 | pathway | 0.5616 | 2.392562 |
| feature:33 | pathway | 0.450318 | 2.20823 | feature:35 | pathway | 0.53299 | 2.270674 |
| feature:54 | pathway | 0.406586 | 1.993779 | feature:33 | pathway | 0.512314 | 2.182589 |

### 3.5 LITERATURE EVIDENCE OF SELECTED PREDICTED ASSOCIATIONS

Based on literature evidence, we attempt to verify the existence of top 20 predicted PPAs, as shown in table 3. For each predicted PPA, we show the mean of the predicted probabilities by XGBoost and lightGBM classifiers, followed by pubmed ids.

**Table 3.** Table showing literature evidence of top 20 predicted HDKP-pathway associations.

| protein-pathway link | average of prediction probabilities | pubmed reference/DOI |
| --- | --- | --- |
| Q9BXA6 and R-HSA-3371511 | 0.999999992 | Pubmed ids:32337545, 38762178 |
| Q86SG6 and R-HSA-3371511 | 0.999999944 | Pubmed id: 40189576 |
| Q9BXA6 and R-HSA-8856825 | 0.999999939 | Pubmed id: 19596796 |
| Q9BXA6 and R-HSA-176187 | 0.999999933 | Pubmed ids: 41959133, 19596796 |
| O94768 and R-HSA-3371511 | 0.999999904 | Pubmed ids: 40953235, 34536317 |
| Q9BXA6 and R-HSA-3371453 | 0.999999874 | Pubmed ids: 21687820, 23599433, 38762178 |
| Q8IU85 and R-HSA-3371511 | 0.999999869 | Pubmed id: 36289281 |
| Q9BXA6 and R-HSA-1679131 | 0.999999866 | Pubmed id: 38762178 |
| Q8N165 and R-HSA-3371511 | 0.999999861 | https://doi.org/10.1016/j.celrep.2021.110233 |
| Q86SG6 and R-HSA-1299308 | 0.999999856 | Pubmed id: 37598857 |
| O94768 and R-HSA-8856825 | 0.999999854 | Pubmed id: 40799387 |
| O94768 and R-HSA-176187 | 0.999999853 | Pubmed id: 40953235 |
| Q9BXA6 and R-HSA-2029482 | 0.99999985 | Pubmed id: 38762178 |
| Q9BXA6 and R-HSA-8948216 | 0.999999849 | Pubmed id: 38762178 |
| Q9BXA6 and R-HSA-3134963 | 0.999999849 | Pubmed ids: 20729278, 38762178, 23599433 |
| Q9Y6M4 and R-HSA-8856825 | 0.999999846 | Pubmed id: 37941124 |
| Q9BXA6 and R-HSA-6785807 | 0.999999845 | Pubmed ids: 19596796, 34831223 |
| Q9BXA6 and R-HSA-5689880 | 0.999999841 | Pubmed ids: 31649732 |
| Q8N2I9 and R-HSA-3371511 | 0.999999835 | Pubmed id: 28089446 |
| Q86SG6 and R-HSA-5689877 | 0.999999827 | Pubmed id: 23973373 |

### 3.6 APPLICATION: PCOS ASSOCIATION AND DRUGGABILITY ANALYSIS

The PCOS relevant HDKPs are enumerated in supplementary table 6. For each HDKP, we consider total pathways associated, total PCOS pathways associated and percentage of associated pathways related to PCOS, sorted in descending order on the latter two parameters.

The HDKPs listed in top ten as per this sorting order and, hence, considered for docking include (UniProt IDs in brackets) PRKACG (P22612), VRK3 (Q8IV63), P1P5K1C (O60331), CSNK2A2 (P19784), PKCθ (Q04759), CAMKK2 (Q96RR4), CSNK1G2 (P78368), MAP3K14 (Q99558), PAK6 (Q9NQU5) and RIO2 (Q9BVS4), having association with 84 to 112 PCOS pathways. These HDKPs are docked with nine popular PCOS drugs namely Alogliptin, Linagliptin, Sitagliptin, Saxagliptin and Vildagliptin (insulin resistance), Letrozole and Clomifene (anovulation), Metformin (insulin resistance and anovulation) and Spironolactone (hyperandrogenism), based on three main issues of treatment viz anovulation, insulin resistance and hyperandrogenism. Docking is performed via MTiAutoDock of MTiOpenScreen webserver [29] which makes use of Lamarckian genetic algorithm in AutoDock Vina 4.2.6 to generate different conformations of drugs. For each drug-protein combination, ten docking runs are conducted and the one with highest binding energy, denoted in kcal/mol is considered. The findings are enumerated in table 4. P22612 shows highest binding affinity with all drugs except Alogliptin. For Alogliptin, the highest binding affinity occurs for Q04759. Among all drugs, the highest binding affinity occurs between P22612 and Spironolactone. Detailed results are available in supplementary table 7.

**Table 4.**
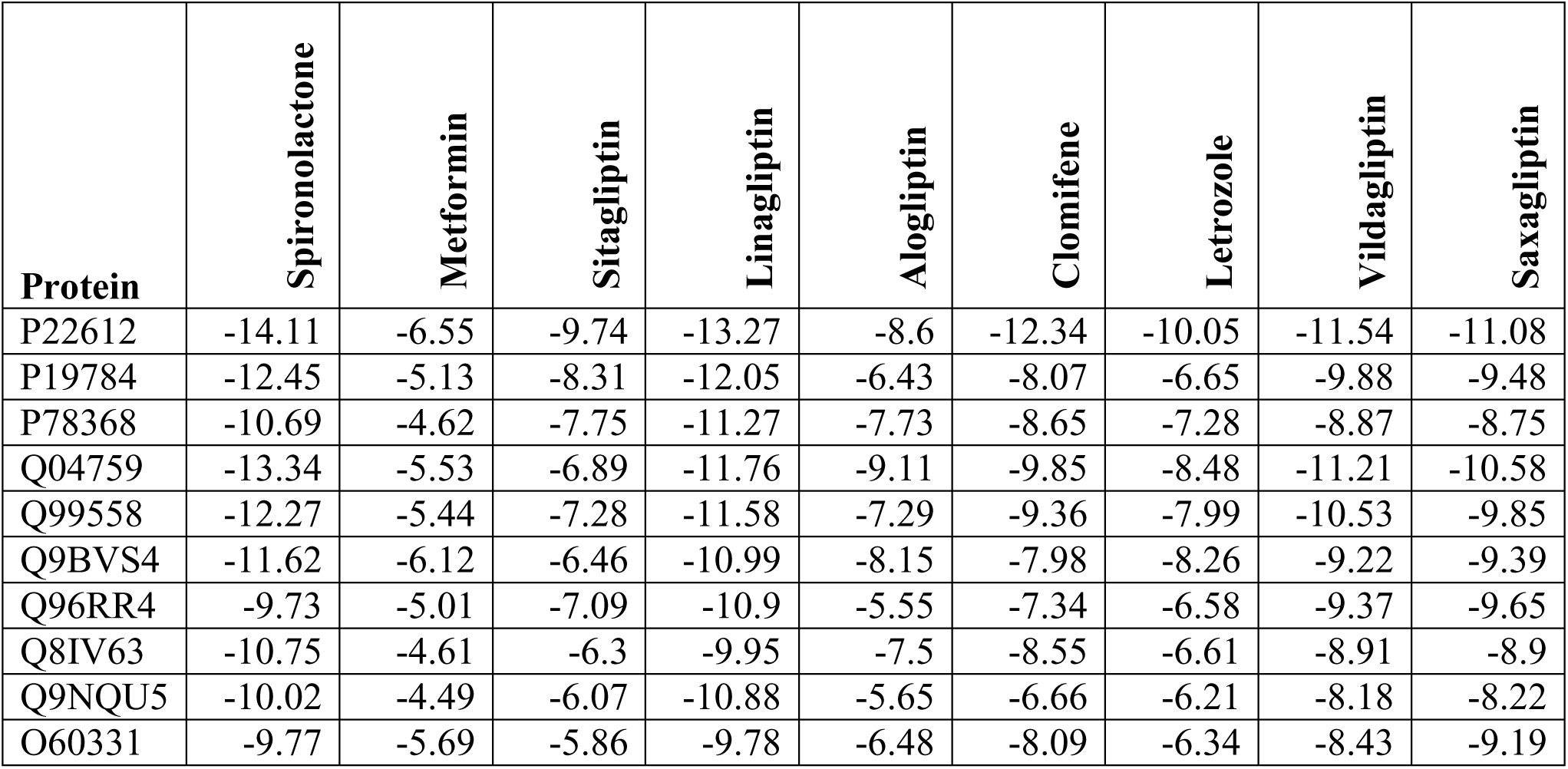
Table showing results of docking experiments conducted on exhaustive combination of nine PCOS drugs and top 10 PCOS-relevant HDKPs, in terms of binding energies released (best poses)

## 4 CONCLUSION

Here, we predict pathway associations of HDKPs by dint of XGBoost and lightGBM classifiers. BERT-based embeddings of proteins and pathways, refined via two-stage transformer and heterogeneous graph transformer and reduced at different stages using PCA and AE along with PPIs and PPAs are used to train both classifiers. Positive (feasible) PPIs and PPAs are curated from databases whereas the negative ones are generated using WGAN-GP. We trace out features responsible for predictions through SHAP-based feature analysis. We infer on functional similarity of HDKPs to human proteins by clustering pathways, mapping each protein to cluster labels of associated pathways (predicted and positive) and conclude on similarity through association to same cluster(s) (denoted by cluster labels). We further trace out PCOS proteins among HDKPs, sort them based on number and percentage of PCOS-associated pathways (predicted and positive) and analyse druggability of a top ten through docking with some known PCOS medications.

We contribute by employing transformer-based language models (BERT-based embeddings) for predicting PPAs involving HDKPs, inferring on functional similarity of HDKPs to human proteins, analysing PCOS-relevance of HDKPs and verifying druggability of a few relevant ones through molecular docking with PCOS medications in vogue. Our model achieves accuracy of 0.9816, precision of 0.9852, recall of 0.9780, f1-score of 0.9816, specificity of 0.9852, MCC of 0.9632, AUROC of 0.9978 and AUPRC of 0.9982, all expressed as mean values over 10 iterations of stratified 10-fold cross-validation. Besides, it correctly classifies 97.48% of test data and predicts 62225 PPAs, all with probability of 0.7 or more. Nine features paramount to and with positive influence on predictions are revealed through SHAP analysis. Ninety six HDKPs are found to have functional similarity to one or more human proteins via clustering. Fifty seven HDKPs are found to be PCOS-relevant. Binding energies in the range of -4.49 to -14.11 kcal/mol are released while docking top ten PCOS-relevant HDKPs with PCOS medications.

In future, our model can be used to derive pathway associations of proteins leading to functionality assessment via involvement in treatment of various diseases.

## ACKNOWLEDGEMENTS

We do hereby acknowledge the use of Supercomputer Paramshakti of the Indian Institute of Technology Kharagpur.

## AUTHOR CONTRIBUTIONS

Somarpita Dutta: Conceptualization, Data curation, Formal Analysis, Investigation, Methodology, Resources, Software, Validation, Visualization, Writing-original draft. Pralay Mitra: Project administration, Supervision, Writing-review and editing.

## FUNDING RESOURCES

This research did not receive any specific grant from funding agencies in the public, commercial, or not-for-profit sectors.

## DECLARATION OF ETHICS

This study did not require ethical approval as it did not involve human or animal participants

